# A role for the VPS retromer in *Brucella* intracellular replication revealed by genome-wide siRNA screening

**DOI:** 10.1101/436246

**Authors:** Alain Casanova, Shyan Huey Low, Maxime Québatte, Jaroslaw Sedzicki, Therese Tschon, Maren Ketterer, Kevin Smith, Mario Emmenlauer, Houchaima Ben-Tekaya, Christoph Dehio

## Abstract

*Brucella,* the causing agent of brucellosis, is a major zoonotic pathogen with worldwide distribution. *Brucella* resides and replicates inside infected host cells in membrane-bound compartments called BCVs (*Brucella-*containing vacuoles). Following uptake, *Brucella* resides in eBCVs (endosomal BCVs) that gradually mature from early to late endosomal features. Through a poorly understood process that is key to the intracellular lifestyle of *Brucella,* the eBCV escapes fusion with lysosomes by transitioning to the rBCV (replicative BCV), a replicative niche directly connected to the endoplasmic reticulum (ER). Despite the notion that this complex intracellular lifestyle must depend on a multitude of host factors, a holistic view on which of these components control *Brucella* cell entry, trafficking and replication is still missing. Here we used a systematic cell-based siRNA knockdown screen in HeLa cells infected with *Brucella abortus* and identified 425 components of the human infectome for *Brucella* infection. These include multiple components of pathways involved in central processes such as cell cycle, actin cytoskeleton dynamics or vesicular trafficking. Using assays for pathogen entry, knockdown complementation and co-localization at single-cell resolution, we identified the requirement of the VPS retromer for *Brucella* to escape the lysosomal degradative pathway and to establish its intracellular replicative niche. We thus validated a component of the VPS retromer as novel host factor critical for *Brucella* intracellular trafficking. Further, our genome-wide data shed light on the interplay between central host processes and the biogenesis of the *Brucella* replicative niche.

**Importance:** With >300,000 new cases of human brucellosis annually, *Brucella* is regarded as one of the most important zoonotic bacterial pathogen worldwide. The causing agent of brucellosis resides inside host cells within vacuoles termed *Brucella* containing vacuoles (BCVs). Although few host components required to escape the degradative lysosomal pathway and to establish the ER-derived replicative BCV (rBCV) have already been identified, the global understanding of this highly coordinated process is still partial and many factors remain unknown. To gain a deeper insight into these fundamental questions we performed a genome-wide RNA interference (RNAi) screen aiming at discovering novel host factors involved in the *Brucella* intracellular cycle. We identified 425 host proteins that contribute to *Brucella* cellular entry, intracellular trafficking, and replication. Together, this study sheds light on previously unknown host pathways required for the *Brucella* infection cycle and highlights the VPS retromer components as critical factors for the establishment of the *Brucella* intracellular replicative niche.

## Introduction

Cellular invasion is a common strategy shared by many bacterial pathogens of human and animals in order to escape host defenses and to establish a protected replicative niche. This notably applies to the human pathogens of the genus *Salmonella*, *Shigella*, *Legionella*, or *Brucella* (1–3). Knowledge of the host cellular pathways that are subverted by these pathogenic bacteria in order to reach and/or establish their intracellular replicative niches can be highly instructive for the development of new treatment strategies. *Brucella* is a facultative intracellular zoonotic pathogen causing animal and human brucellosis. With more than 300,000 new cases of human brucellosis every year, *Brucella* is regarded as one of the most important zoonotic bacterial pathogen worldwide (4–6). There is currently no effective vaccination for humans and even prolonged combinatory antibiotic treatments do not fully protect against relapses (7). Therefore, *Brucella* remains a significant threat to public health and to the economy in endemic areas, and thus new treatment strategies to circumvent *Brucella* infections are highly needed.

At the cellular level, *Brucella* invades both phagocytic and non-phagocytic cells where bacteria persist and replicate inside membrane-bound compartments – the *Brucella* containing vacuoles (BCVs). BCVs sequentially interact with components of the host early and late endocytic pathway (eBCVs) then transit to establish the replicative niche (rBCVs) in vesicles that are directly connected to the endoplasmic reticulum (ER) (8) and harbor ER-specific markers, such as SEC61 and calnexin (8–10). Several critical steps for the intracellular journey of *Brucella* and their associated host factors have been resolved. For instance, adherence to the host cell surface is mediated via interaction with sialic acid residues or binding to fibronectin and vitronectin (11, 12). Internalization requires actin remodeling via the activity of the small GTPases RAC, RHO and direct activation of CDC42 (13). Upon internalization, bacteria are contained within eBCVs that successively associate with a subset of endosomal markers, starting with RAB5, the early endosomal antigen (EEA1), the transferrin receptor (TfR), as well as the lipid rafts component flotillin-1 (9, 14-16). Next, the eBCVs associate with the late endosomal markers RAB7, RILP (RAB7’s effector RAB interacting lysosomal protein), LAMP-1 (Lysosomal-associated membrane protein 1), and transiently with autophagosomal markers (9, 10). As they evolve late endosomal characteristics, acidification of the eBCVs serves as a trigger for the expression of the VirB type IV secretion system (T4SS, (17, 18)). This major pathogenicity factor is required to prevent the complete fusion of eBCVs with lysosomes, consequently allowing a fraction of the internalized *Brucella* to avoid host-mediated degradation, and promotes the maturation of the eBCVs towards the rBCVs (16, 18). Noteworthy, the T4SS effectors responsible for this escape remain largely elusive, despite a growing repertoire of identified candidates (recently reviewed in (19)). Most recently, it was discovered that subversion of both anterograde and retrograde transport and recruitment of Conserved Oligomeric Golgi (COG) tethering complex-dependent vesicles to the BCV promotes the establishment of the *Brucella* replicative niche (20). Importantly, despite all these findings, the precise mechanism(s) resulting in diversion of eBCVs from the endolysosomal pathway towards the ER-associated replicative compartment (rBCVs) is still largely unresolved. The same holds true for host factors required for maintenance of the replicative niche.

In this study, we took a systems-level approach to gain a deeper insight into these fundamental questions. Using a genome-wide RNA interference (RNAi) screening approach, we identified 425 host proteins whose knockdown either increases (202) or decreases (223) *Brucella* intracellular replication. Beside the rediscovery of several previously identified host targets, that validates our approach, data reveals numerous novel candidate components that can modulate *Brucella* cellular entry, trafficking, and/or replication. Among these targets, we identified VPS35 and VPS26A, two components of the trimeric <u>v</u>acuolar <u>p</u>rotein <u>s</u>orting (VPS) complex (termed here VPS retromer), which are required for the diversion of BCVs from the endolysosomal pathway and the establishment of the intracellular replicative niche.

## Results

### A genome-wide siRNA screen reveals novel host pathways involved in *Brucella* infection

To identify novel host factors important for *Brucella* intracellular infection, we performed a genome-wide siRNA perturbation screen on the human epithelial cell line HeLa (ATCC© CCL-2) combined with bacterial infection at biosafety-level 3. Infections of the siRNA-treated cells were performed with a GFP expressing strain of *B. abortus* and the outcome was analyzed at 48 h post-infection (hpi) using automated fluorescence microscopy (21, 22). Infection scoring was determined with a tailored high-content analysis workflow (Fig. 1 and Materials and Methods). In brief, a model of *Brucella* replication was fitted to the pathogen intensity distribution to gain an infection classification independent of absolute fluorescence intensity. Further, we implemented an image intensity normalization coupled with a novel pathogen-to-cell association approach, which enabled quantitative measurement of the pathogen intensity distribution (Materials and Methods). An overview of the results is presented in Fig. S1. To account for the well-known confounding off-target effects associated with siRNA technology (recently reviewed in (23)), we applied a multiple orthologous RNAi reagents approach (MORR (24)) with n ≥ 5 perturbations per host gene. Further, we applied the Redundant siRNA Analysis (RSA) algorithm (25) on the entire screening data to reduce the number of false positives caused by off-target effects of single siRNAs and to favors genes with a reproducible phenotype confirmed by independent siRNAs. Genes matching a Benjamini-corrected RSA p-value ≤ 0.01 with more than 3 hit wells were considered as significant and selected for further analysis (see also Material and Methods). As a result, we identified 425 significant hits affecting *Brucella* infection. These comprised 223 down-hits (Fig. S1A, red and Table S1) and 202 up-hits (Fig. S1A, green and Table S2). Single siRNA data points are presented in Fig. S2 (down-hits) and Fig. S3 (up-hits). A panel of representative images from the screen is presented in Fig. 2A. Hit genes were further stratified by gene-annotation enrichment analysis and functional annotation clustering using DAVID (26), protein-protein interaction network using the STRING database (27), as well as manual datamining. The functional categories enriched in our hit lists are presented in Fig. 2B-D together with the high confidence protein-protein interaction network for targets that reduced (Fig. 2E) or increased (Fig. 2F) *Brucella* infection upon knockdown. Gene ontology and functional clustering analysis indicated a rather small overlap in enriched pathways when considering up or down hits (Fig. 2B-D). The most prominent clusters that positively affected infection upon knockdown comprised components involved in the control or the modulation of central cellular processes such as protein synthesis, transcription and mRNA processing, and cell cycle progression, as well as clathrin-mediated endocytosis (Fig. 2 and Table S2). The most prominent clusters that negatively affected infection upon knockdown comprised signaling pathways involved in actin-remodeling and phagocytosis. These included most core components of the Actin-related protein-2/3 complex (ARP2/3: ARPC2, ARPC3, ACTR2 and ACTR3), and one of its main modulator, the WASP regulatory complex (WRC: CYFIP1, WASF3, NCKAP1, and ABI3). Down-hits also comprised multiple components involved in TGF-β and Eph signaling as well as further vesicular/endocytic pathways (Fig. 2 and Table S1). Among all these factors we can highlight the Ras related protein RAB7A, which is needed for *Brucella* trafficking to the replicative niche (10), the small GTPases RAC1 and CDC42, which are involved in *Brucella* internalization into non-phagocytic cells (13) as well as the transmembrane glycoprotein SLC3A2 (CD98hc), involved in both bacterial uptake and intracellular multiplication (28). Since the role of these individual components has already been described in the context of *Brucella* infection, they can be considered as benchmark to our results, and globally validate our systems-level perspective of the human infectome for *Brucella* infection.

**Fig. 1.**
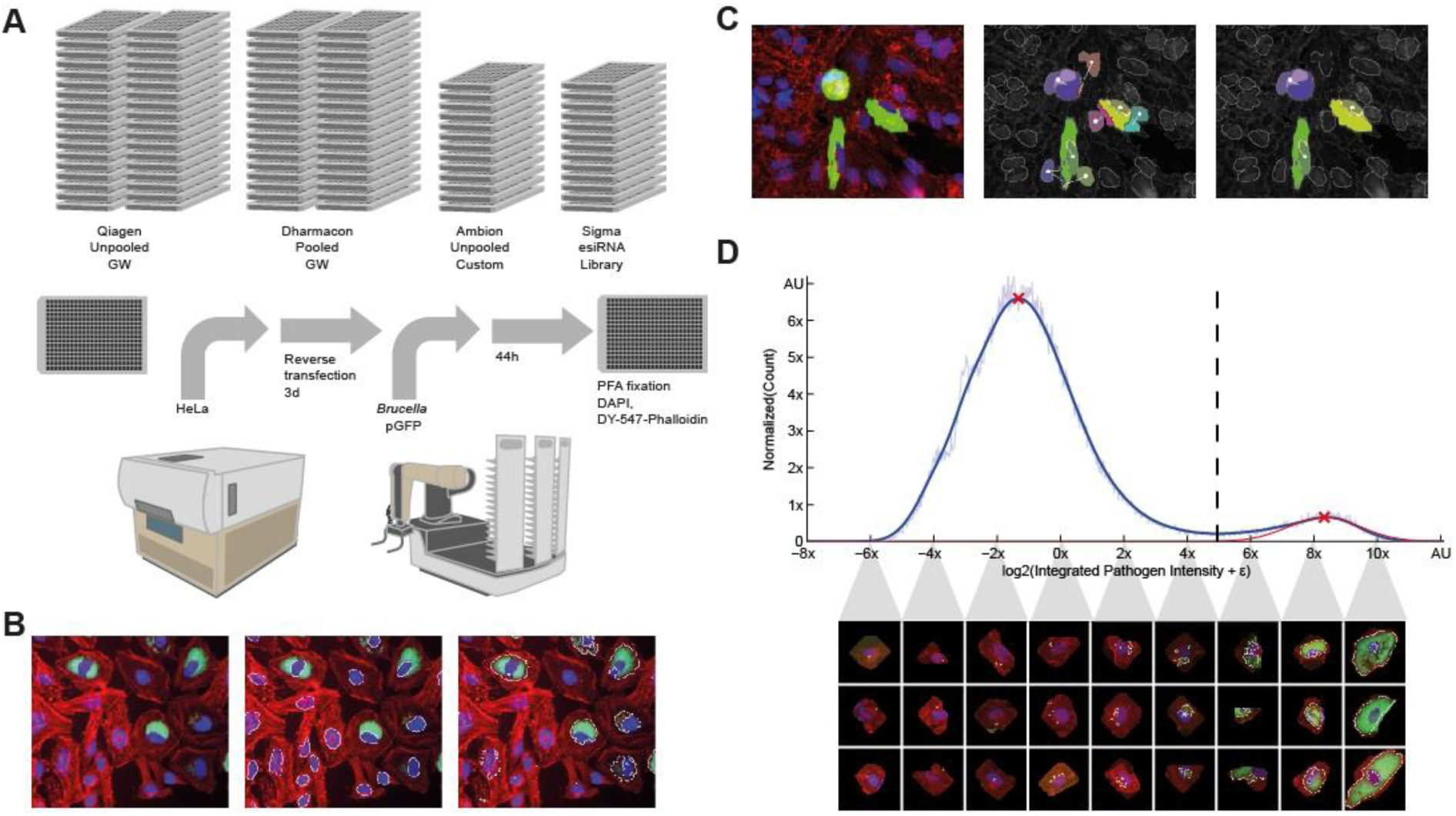
**Overview of the high-content screening and analysis**. (**A**) Summary of RNAi screening workflow. Reverse transfection of HeLa cells was performed in 384-well format for 72 h followed by 48 h infection with GFP expressing *B. abortus*, PFA fixation, and staining of HeLa cells with DAPI and DY-547-Phalloidin before automated imaging. (**B**) Image analysis was performed with CellProfiler to segment nuclei and bacteria and to extract measurements. (**C**) Accurate association of segmented bacteria to nuclei enables quantitative single cell measurements. The naive association (middle image) of segmented pathogen can be affected by over-splitting in dense cell populations (left image). Our proposed solution (right image) based on a nucleus attraction score. (**D**) The plate histogram shows the bimodal distribution of integrated GFP intensity corresponding to *Brucella* replication. Intensity on the X-axes is log2-scaled to account for exponential growth. The normal distribution fitted (red curve) to the Kernel Density Estimation of the histogram allows to compute a robust binary infection threshold (dashed line) separating HeLa cells with (right) and without (left) replicating *Brucella*. Associated are samples of single-cell images corresponding to the intervals of the intensity distribution (for more details see Materials and Methods).

**Fig. 2.**
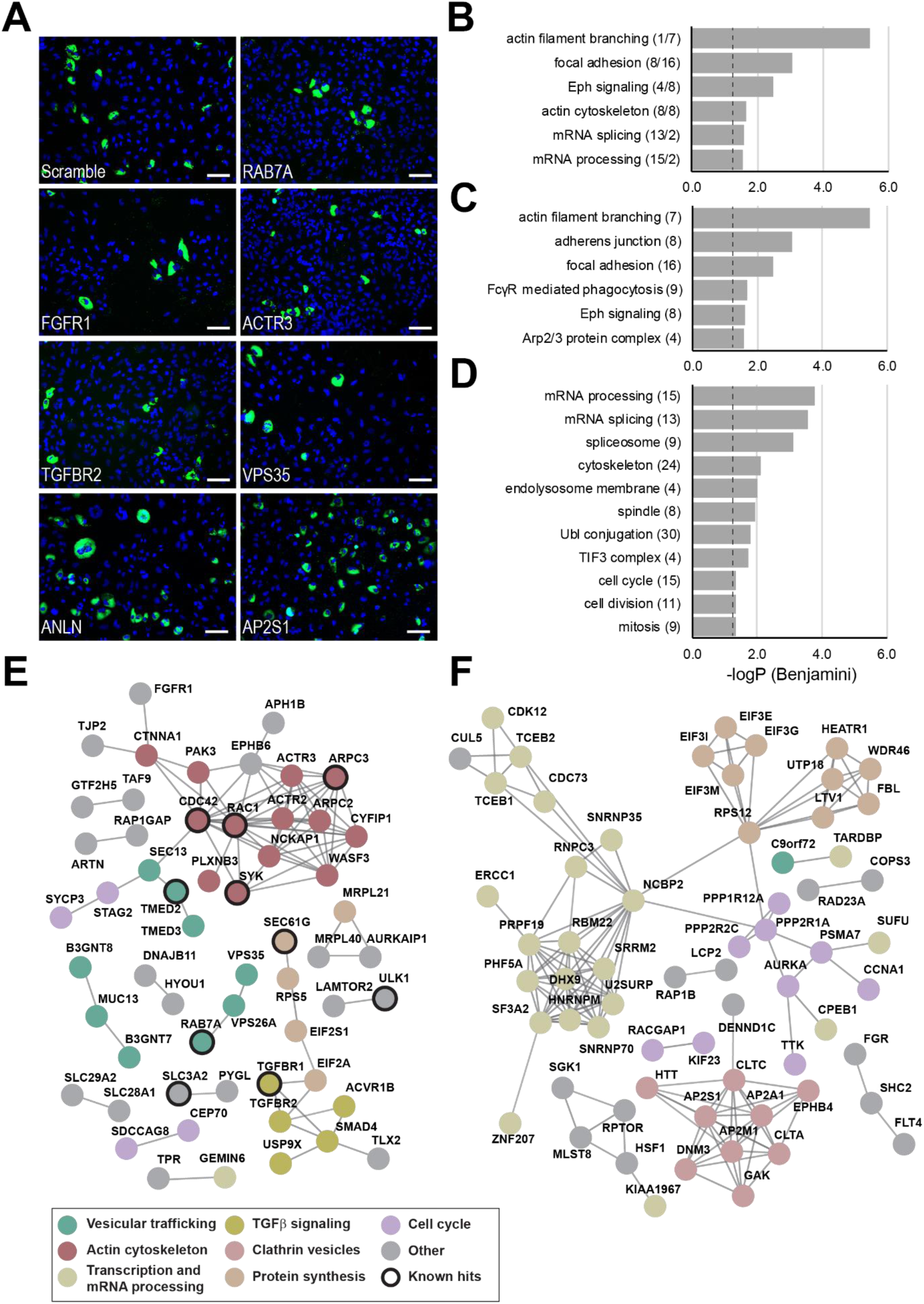
The *human* infectome for *Brucella* infection determined by genome-wide siRNA screening. **(A)** Representative merged images from the genome wide screen showing nuclei (DAPI) and intracellular replication of GFP expressing *Brucella abortus* for either control conditions (scramble) or a panel of identified hits (RAB7A, FGFR1, ACTR3, TGFBR2, VPS35, ANLN and AP2S1). Scale bar = 100 µm. **(B, C, D)** Results of gene ontology enrichment analysis (DAVID) for the entire hit list (B), the down- (C) or up-hits (D). The [-log_10_] of the P-value associated to the different categories are indicated (cut-off: Benjamini corrected P-value ≤ 0.01, with more than 3 hit wells) as well as the number of individual components associated to the displayed categories are indicated. In (B), the first number refers to down-hits and the second to up-hits. **(E, F)** High confidence protein-protein interaction networks (x ≥ 0.8) for the 223 RSA down-(A) or 202 up-hits (B) determined using the STRING database. Clusters with common predicted cellular function are colored and their prominent function are indicated. Hits previously reported to be involved in *Brucella* pathogenicity are highlighted: ARPC3 (29); TGFBR2 (22); SEC61G and TMED2 (35); RAC1 and CDC42 (13); SLC3A2 (28); RAB7A (10); SYK (63); ULK1 (64); For clarity, disconnected components are not displayed. The complete list of down- and up-hits is presented in Tables S1-S2. Data from single RNAi reagents for down- and up-hits is presented in Fig S2-S3.

### Pathogen entry assay identifies a role for VPS35, VPS26A and SEC61γ in *Brucella* post-entry trafficking

To further dissect the role of the identified genes in the progression of *Brucella* infection, we took advantage of a pathogen entry assay previously developed in our laboratory (29). Briefly, at 4 hpi host cell membrane-impermeable gentamicin was added to selectively kill extracellular *Brucella* and concomitantly cell membrane-permeable anhydrotetracyclin was added to induce expression of a plasmid-encoded reporter in the viable intracellular bacteria. At 8 hpi this approach allowed us to robustly identify individual intracellular bacteria and to quantify the bacterial load before intracellular replication is initiated (Fig. 3A and Material and Methods). For this assay, we selected a number of genes from the different pathways identified in the genome-wide screen as well as additional genes supplementing them (Table S3). The results of this entry assay were plotted against a matching endpoint assay (intracellular replication at 48 hpi). Strikingly, most of the tested genes displayed a direct correlation between the results of the entry and the endpoint assay (r^2^ =0.763). This was the case for components involved in the actin-remodeling pathway (RAC1, ACTR3, CYFIP1) or those involved in the TGFβ signaling (SMAD4, TGFBR1, TGFBR2), which strongly reduced both entry and subsequent intracellular replication (Fig. 3B and Table S3). Similarly, the components of the clathrin pathway GAK and AP2S1 both increased bacterial entry and replication (Fig. 3B and Tables S3-4). This support their involvement in *Brucella* entry into non-phagocytic cells, without excluding an additional role at any further stage of the infection. To identify components with a divergent outcome between entry and replication, we selected genes diverging by more than one standard deviation to the fitted data. Six genes matched this criterion (Fig. 3B). Three genes displayed an apparent higher effect on pathogen entry than subsequent replication (albeit at a rather modest level). These were the small GTPases CDC42 (13), the α1 subunit of the Na+,K+-ATPase ATP1A1(30) and the subtilisin-like endoproteinases FURIN (31). Most strikingly, three genes displayed a stronger reduction in endpoint replication compared to entry. Knockdown of Sec61γ - a central element of the ER-protein translocation machinery (see for instance (34), which has been previously involved in *Brucella* infection (35), showed a strong decrease in intracellular replication albeit no effect on pathogen entry. Similarly, our assay identified the <u>v</u>acuolar <u>p</u>rotein <u>s</u>orting associated proteins VPS35 and VPS26A - two essential components of the VPS retromer complex (recently reviewed in (32, 33)). These genes and associated pathway(s) thus likely represent novel components controlling the post-entry trafficking of *Brucella* towards its replicative niche and/or are themselves required for the establishment or maintenance of the rBCVs. For the present study, we further focused on the role of VPS35 and the VPS retromer in *Brucella* trafficking as it was the most prominent hit in our entry assay.

**Fig. 3.**
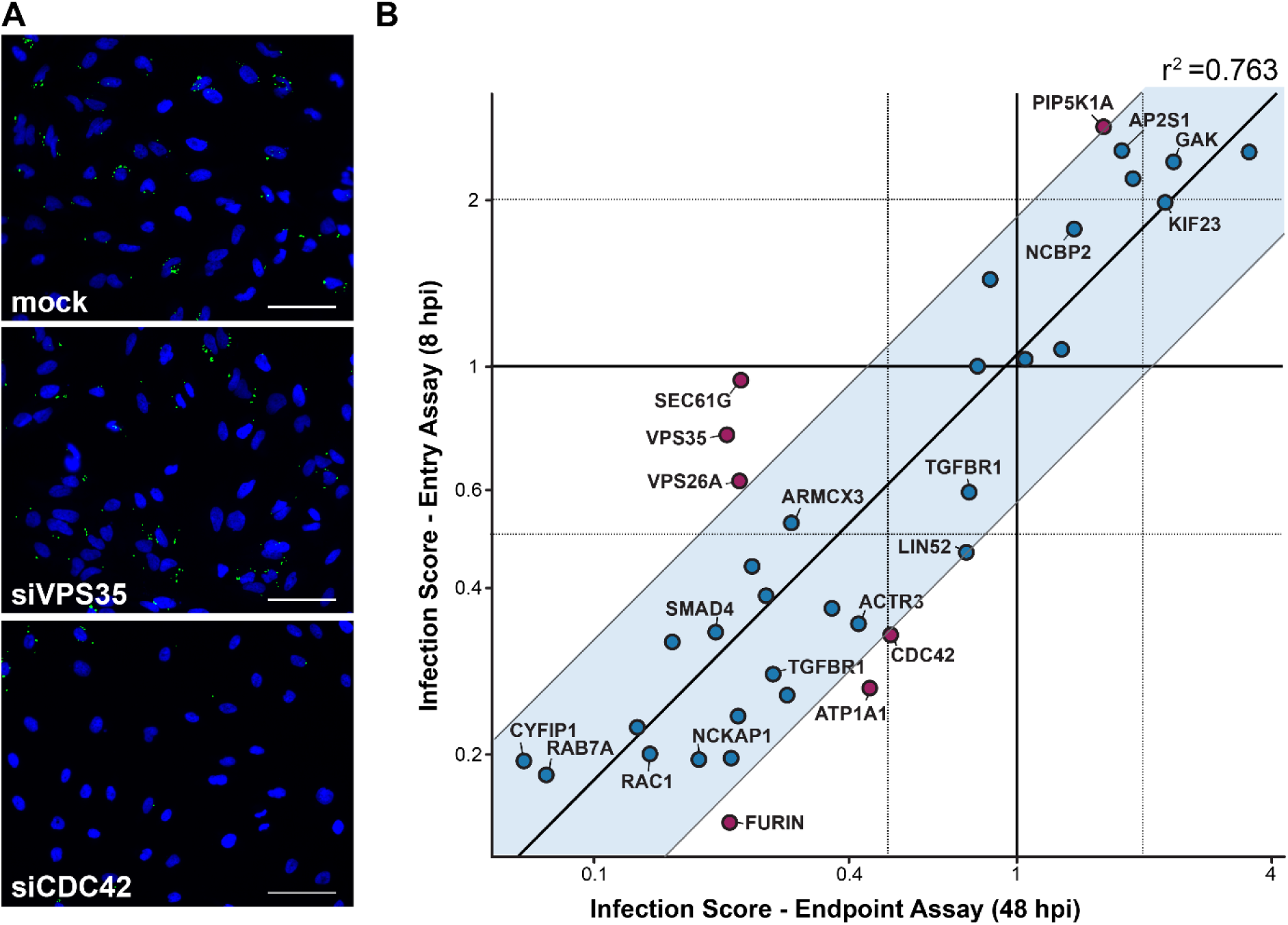
Entry assay identifies new components required for post-entry processes during *Brucella* infection. **(A)** Representative images from the entry assay showing nuclei (DAPI) of HeLa cells and intracellular *Brucella abortus* (GFP) for control condition (mock) and cells treated with siRNAs against CDC42 or VPS35. Scale bar = 100 µm. HeLa cells were infected with *B. abortus* expressing GFP under a tetracycline inducible system for 8 h (see Material and Methods). **(B)** Scatter plot in double logarithmic scale showing infection scores measured for the entry assay (8 hpi) versus endpoint assay (48 hpi), normalized to the respective mock dataset (Table S3). For the entry assay, cells containing single bacteria were considered as infected and the final readout is proportional to the median number of bacteria per infected cells. For the endpoint assay, only cells containing replicating bacteria were considered as infected (Fig. 1 and Materials and Methods). Each data point corresponds to the average of all siRNAs or esiRNAs targeted against the gene of interest (n=3). The straight fit (oblique line, r^2^ = 0.763) indicates a globally high correlation between both assays. The blue box comprise all points within ± 1 SD to the fitted data. The genes falling out of this range are marked in red. For ease of visualization, only the averaged values over all RNAi products targeting a given gene are displayed.

### The VPS retromer is important for *Brucella* intracellular replication

The retromer complex orchestrates the recycling of numerous transmembrane proteins from early and maturing endosomes either to the trans-Golgi network (TGN) or back to the plasma membrane. Formed by a heterotrimeric complex consisting of VPS26, VPS29, and VPS35, the VPS retromer is conserved from yeast to human. However, the individual retromer sub-complexes have functionally diverged to organize multiple distinct sorting pathways, depending on the association with different accessory factors (32, 33). To further decipher the role of the retromer in *Brucella* trafficking and intracellular replication we specifically browsed our genome-wide siRNA data for retromer-associated proteins (Fig. 4A and D). Further to VPS35 and VPS26A, already identified both in the genome wide and in the entry screen (Fig. 2E and Fig. 3B), knockdown of VPS26B (the paralogue of VPS26A) resulted in a significant reduction in *Brucella* infection (Fig. 4D). Depletion of VPS29, the third core component of the VPS retromer, resulted only in a mild reduction of *Brucella* infection and did not reach significance due to the wide spread of data obtained for the cohort of 9 individual siRNAs tested (suggestive of strong off-target effects). Next to the retromer component, knockdown of the small GTPase RAB7A showed the strongest reduction in intracellular *Brucella* (Fig 4D). However, neither SNX3, that together with the VPS retromer forms the SNX3 retromer nor SNX27, another retromer-associated component involved in endosome-to-plasma membrane trafficking (36, 37), displayed significant effect (Fig. 4D). Depletion of SNX1 and SNX5, two of the four sorting nexins of the SNX-BAR retromer (38), even seems to enhance *Brucella* infection (although they did not pass our hit-selection criterion) while the two others, SNX2 and SNX6, showed no effect (Fig. 4D). Noteworthy, the functional association of the SNX-BAR sorting nexins with the VPS retromer has been challenged by two recent publications, which rather support a VPS-independent action of SNX-BAR (39, 40). Collectively, our data indicate that the observed post-entry impairment in *Brucella* intracellular replication is specifically linked to the integrity of the heterotrimeric VPS retromer, although involvement of further components in this process cannot be excluded.

**Fig. 4.**
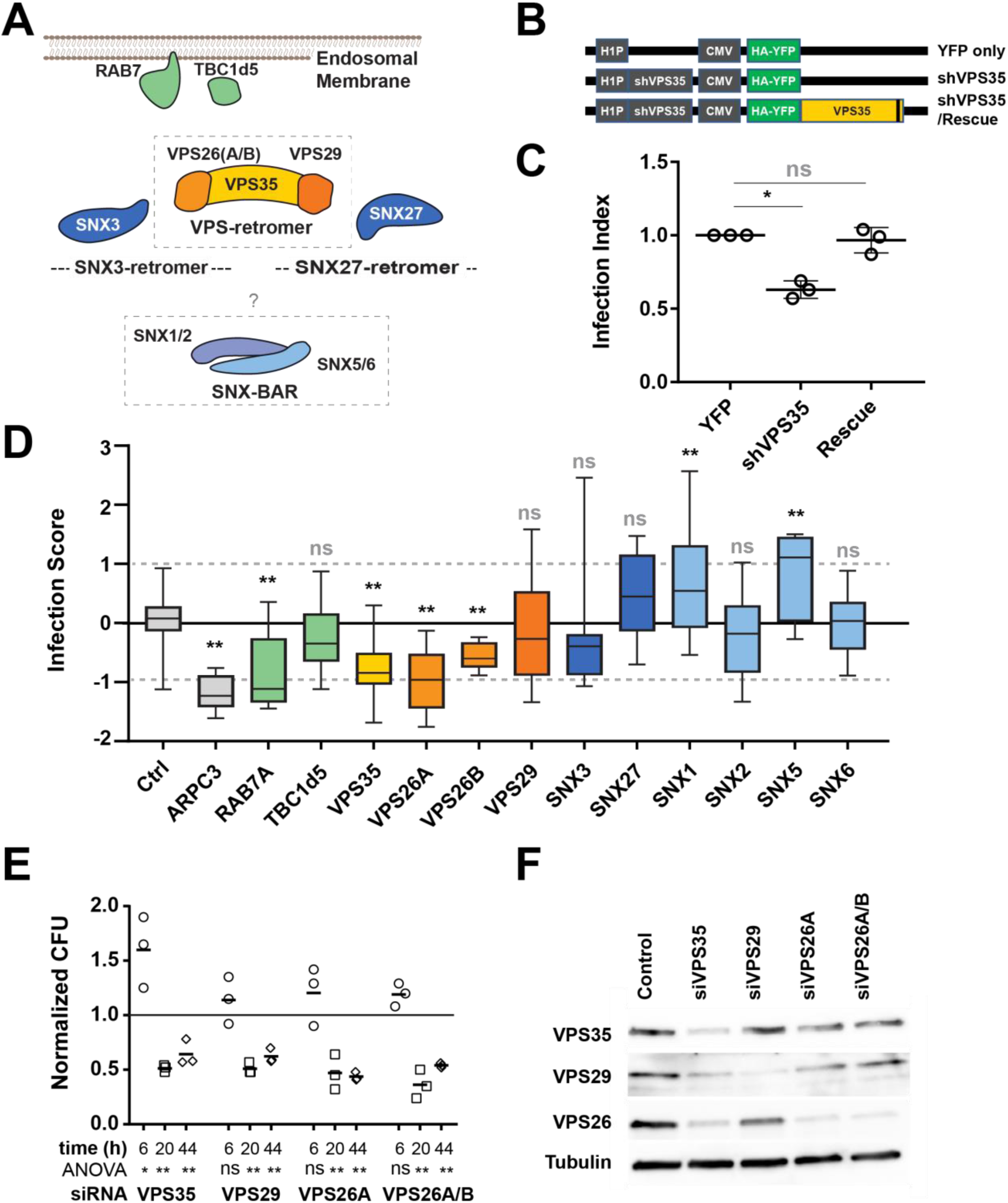
The VPS retromer is a key component of *Brucella* intracellular trafficking. **(A)** Schematic representation of the retromer components and their prominent interactors. (**B**) Schematic representation of the shRNA constructs used in (C). The gray box on the shVPS35 / Rescue construct indicates the silent mutations that prevent recognition by to the co-expressed shRNA (41). (**C**) Infection index from transfected cells. Displayed are the averaged infection index and associated standard deviation after 48 h of *Brucella* infection. Data was normalized to the YFP only condition (n=3). Asterisks indicate statistically significant difference to scrambled YPF only condition as determined by paired t-test (* Pval ≤ 0.01, ns not significant). (**C**) Dot box representation of the z-scored infection score for components of the retromer and interactors, including the positive control ARPC3. Asterisks indicate statistically significant difference to scrambled siRNA-treated bacteria (Ctrl) as determined by one-way ANOVA and Dunnett’s multiple comparisons test (** Pval ≤ 0.001; ns not significant). (**E**) Normalized colony forming units (CFU) recovered from siRNA-treated cells a 6, 20 or 44 hpi. The presented data correspond to CFU count normalized to control-siRNA-treated cells (n=3). Significance was determined using One-way ANOVA with Dunnet’s multiple comparison test (* Pval ≤ 0.01; ** ≤ 0.001; ns not significant). (**F**) Western blot analysis of the indicated proteins in total lysate of Hela cells treated with siRNA targeting the designated genes, 72 h post transfection. Displayed is a representative example of an experiment performed in biological triplicate (n=3). See Table S4 for the matching averaged intensity quantification.

To validate the requirement of VPS35 on *Brucella* infection and to rule out any off-target effects, we performed a complementation experiment using a VPS35 cDNA insensitive to a co-expressed shRNA (41). While shRNA knockdown of endogenous VPS35 inhibited *Brucella* infection, as detected in our genome-wide approach, ectopic expression of the shRNA-insensitive cDNA of VPS35 rescued the phenotype (Fig. 4B and C), confirming that depletion of VPS35 indeed negatively affects *Brucella* infection. We further confirmed the observed requirement of the VPS retromer for *Brucella* intracellular replication by determining intracellular bacterial load at different infection time of siRNA-transfected cells using colony forming unit (CFU) determination (Material and Methods). At 6 hpi, no significant difference to the control was detected, with the exception of a small increase in intracellular bacteria upon VPS35 knockdown (Fig. 4E). Importantly, at 20 h and 44 hpi, siRNA knockdown of either VPS35, VPS29, or VPS26 resulted in a significant decrease of viable intracellular *Brucella* compared to control-treated cells (Fig. 4E) confirming the data obtained by our microscopy-based entry screen (Fig. 3). Further, efficiency of siRNA knockdown was confirmed by Western blot analysis (Fig. 4F and Table S4). Together, these results corroborate the importance of each constituent of the VPS retromer, including VPS29, for *Brucella* to reach and possibly to maintain its intracellular replicative niche.

### VPS35 knockdown prevents *Brucella* escape from the lysosomal pathway

Transient association with the lysosomal marker LAMP-1 is a hallmark of BCV trafficking during the first hours of infection. This association is eventually lost for those bacteria that manage to escape the host degradative pathway. Thus, to investigate the role of VPS35 in *Brucella* trafficking and to assess at which stage it could be required for the establishment of the intracellular replicative niche, we quantified *Brucella* co-localization with LAMP-1 in siRNA-treated and control cells. To this end, we analyzed *Brucella*-infected HeLa cells at 6 and 18 hpi and determined the percentage of LAMP-1 co-localization for each detected bacteria, combining immunostaining and confocal microscopy (Fig. 5 and Materials and Methods). At 6 hpi, most *Brucella* were found within LAMP-1-positive vesicles in both control and siRNA-treated cells (Fig. 5A-D, F), indicating that VPS35 function is not required for the early trafficking of the BCVs. However, loss of LAMP-1 association at 18 hpi was mainly detected in control cells whereas most *Brucella* remained in a LAMP-1 positive compartment upon VPS35 knockdown (Fig 5B, C). Accordingly, VPS35 depletion strongly reduced intracellular replication of *Brucella* 18 hpi (Fig. 5G) compared to control cells (Fig. 5E).

**Fig. 5.**
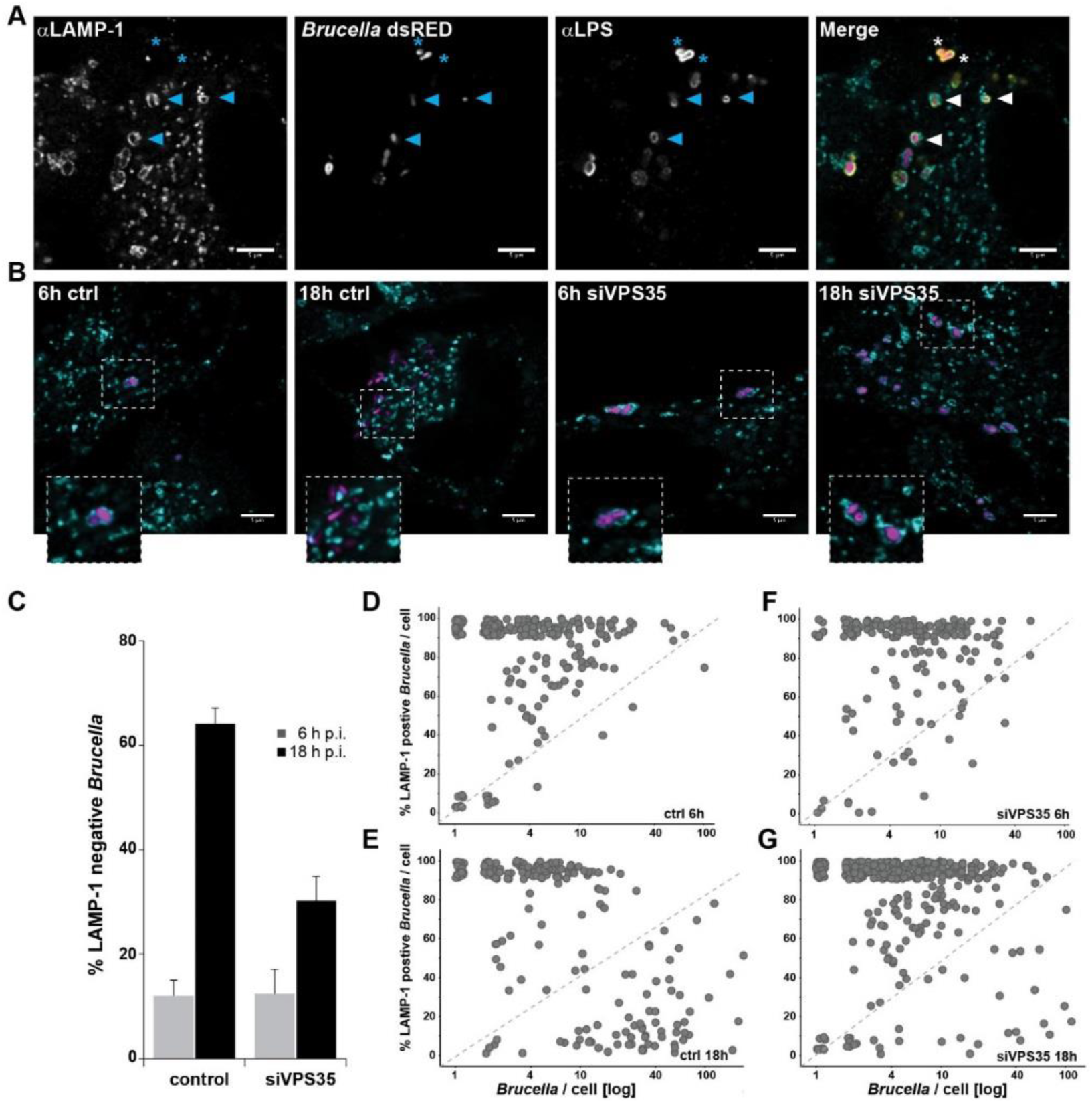
VPS35 is required for *Brucella* to escape the lysosomal pathway. (**A**) Immunofluorescence approach used to quantify localization of *Brucella* within LAMP-1 positive vesicles, illustrated with a representative example of control-treated cells 6 hpi. Individual channels and merged picture are presented. Arrows indicate examples of co-localization of bacteria with LAMP-1 positive compartments. Staining of *Brucella* LPS (αLPS) was used to confirm the presence of LAMP-1 in direct proximity of the bacterial surface. Asterisks indicate examples of LAMP-1 negative *Brucella*. (**B**) Representative images from *Brucella* infected cells either mock transfected (ctrl) or after VPS35 knockdown (siVPS35). Samples were fixed 6 and 18 hpi. For clarity only the LAMP-1 (cyan) and dsRed (magenta) channels are presented. Scale bar: 5 μm. (**C**) Global quantification of LAMP-1 negative *Brucella.* Displayed are the average and associated standard deviation for more than 500 bacteria and more than 50 Hela cells per time point and condition (n=3). (**D**) Single-cell data representation of the data presented in (C). Displayed is the distribution of LAMP-1 positive *Brucella* per cell as a function of the total number of bacteria counted in that given cell.

Altogether, our single cell co-localization analysis demonstrates the requirement of VPS35, and thus of a functional VPS retromer, for the diversion of BCVs from the lysosomal pathway and for the subsequent establishment of a successful replicative niche.

## Discussion

The different membrane-bound organelles that compose the secretory pathway and the endo-lysosomal system of eukaryotic cells constitute targets of choice for many intracellular pathogens, which have evolved highly diverse strategies to hijack and/or subvert these trafficking pathways to their benefit (2, 3). In that context, the importance of retrograde trafficking for the infection cycle of a number of human pathogens (viruses and bacteria) has been recognized in the past years (e.g., (42, 43)). This is for instance the case for *Chlamydia trachomatis*, which uses its effector LncE to subvert host restriction via direct interaction with SNX5, thereby disrupting retromer trafficking (44, 45). Further pathogens have been shown to specifically target the VPS retromer, using or subverting its function to their advantage. For instance, the Hepatitis C virus interacts with VPS35 through its protein NS5A. This viral protein is recognized as VPS retromer cargo and its interaction with VPS35 supports viral replication (46). Among bacterial pathogens, the best-studied example to date is the subversion of the VPS retromer function by the T4SS effector RidL of *Legionella pneumophila.* RidL was shown to interact with VPS29, inhibiting retromer activity by outcompeting the binding of the VPS retromer regulator TBC1d5 and thereby promoting *Legionella* intracellular replication (47–49). Most recently, integrity of the *Salmonella* containing vacuole was shown to be maintained by the direct interaction of the SPI-2 T3SS effector SseC with the VPS retromer (50). In this study, we report the involvement of the VPS retromer in *Brucella* intracellular cycle. More specifically, we show that the VPS retromer integrity is required for *Brucella* to escape the host degradative pathway, as supported by the inability of the eBCVs to mature upon VPS35 knockdown towards LAMP-1 negative rBCVs. We further show that this VPS-retromer-dependent process takes place after internalization and early trafficking, between 6 h and 18 hpi, matching the estimated timing of the eBCV-to-rBCV transition. It is thus tempting to speculate that the VPS retromer, possibly together with RAB7A, plays a role in this yet elusive but essential branching point of *Brucella* intracellular trafficking i.e. diversion from the lysosomal pathway towards its ER-associated replicative niche. An alternative, yet non-exclusive hypothesis is that the VPS retromer is involved in the establishment of the rBCV, possibly by providing host factors and/or membranes that follow retrograde trafficking. A further role for the VPS retromer in the maintenance of the rBCV cannot be excluded based on our results. With the accumulation of functional data and the increasing number of described interactors, the VPS retromer is nowadays largely appreciated as recruiting hub that orchestrates the retrograde endosomal trafficking of numerous cargos to the TGN or the plasma membrane (32, 33, 51). This versatility however obscures the identification of the underlying mechanism(s) by which VPS35 and the VPS retromer may contribute to *Brucella* intracellular fate. Further browsing our dataset for the effect of known VPS retromer interactors failed resolving the VPS retromer-dependent pathway(s) – if any – that is required for *Brucella* intracellular replication. The only VPS retromer interactor that we unambiguously identified is the small GTPase RAB7A, which is essential for the recruitment of the retromer to endosomal membranes (52, 53). Importantly, association of this upstream interactor to the eBCV is a well-established hallmark of early *Brucella* intracellular trafficking (9, 10, 16). Moreover, over-expression of a RAB7 dominant negative allele (RAB7^T22N^) impairs the establishment of the *Brucella* replicative niche (10). This finding was recapitulated by our siRNA knockdown approach, strengthening the role of RAB7 in controlling BCVs’ fate, albeit by an unknown mechanism. Considering that the recruitment of the retromer to endosomal membranes is strictly dependent on the presence of RAB7, it is conceivable to assume that depletion of RAB7 prevents the recruitment of VPS35 to the BCV, consequently explaining the RAB7-dependency observed for *Brucella* replication. Alternatively, very recent findings have established that retromer depletion in Hela cells actually results in the hyper-activation of RAB7, which causes an overall depletion of the RAB7 pool on endo-membranes (54). That drastic consequence could also imply an indirect effect of the observed retromer requirement for *Brucella* trafficking, by acting at the level of RAB7 activity and its availability for the BCV maturation. However, our results indicate that the effect of RAB7A or VPS35 siRNA knockdown are not entirely congruent. Whereas depletion of either factor impairs *Brucella* intracellular replication, only RAB7A knockdown showed a marked effect on pathogen entry whereas VPS35 appears to be only required at a later stage of the infection. The implications of the newly described feedback signaling on RAB7 triggered by the retromer depletion, as well as the relative contribution of the VPS retromer and RAB7 for the transition of the eBCVs to rBCVs should be addressed in future studies.

Besides the retromer complex, our study pinpointed the involvement of several host pathways in *Brucella* infection, which had not yet been associated with this process. The most prominent cluster negatively affecting infection upon knockdown comprises factors involved in actin remodeling and actin dynamics as well as associated signaling pathways. Apart from an early association of Arp2/3 with BCVs (55), surprisingly little is known about the exact role played by the Arp2/3 complex or the WASP regulatory complex and their regulators during and possibly after *Brucella* internalization. Here as well, further studies will be needed to decipher the precise nature of their involvement. Finally, we also found that members of the TGF-β and FGF signaling pathways promote *Brucella* infection as their depletion resulted in decreased *Brucella* infection. Interestingly, it has previously been reported that patients with brucellosis show higher TGF-β1 serum levels, a finding that is correlated with depressed T cell function (56). Further, B cells were also shown to produce TGF-β at early stages of infection with *Brucella* in mice (57). A possible immunosuppressive role for this pathway during *Brucella* infection should be further investigated.

Summing-up, we believe that the genes and pathways identified in this study constitute a rich resource towards the understanding of *Brucella* intracellular trafficking, which ultimately should allow development of new approaches to controlling *Brucella* infections in human.

## Material & Methods

### Cell lines and plasmid constructs

All experiments were performed in the human cervical carcinoma epithelial cell line (Hela) ATCC© CCL-2. Infections were performed using *Brucella abortus* 2308 carrying the constitutive GFP expression plasmid pJC43 (*aphT::GFP* (58)), pAC042.08 for entry assay (*apht::dsRed,tetO::tetR-GFP* (29)) or pAC037 (*apht::cerulean*, this study) for rescue experiments. Cells and bacteria were grown as described in (22, 29). pAC037 was constructed by replacing dsRed from pJC44 (10) with Cerulean from pCERC-1 (59). Cerulean was amplified using prAC082 (TGGATCCGAAAGGAGGTTTATTAAATGGTGAGCAAGGGCGAGGAGC) and prAC083 (TCTAGAGCTAGCTTACTTGTACAGCTCGTC) and cloned into pJC44 by restriction/ligation using BamHI and XbaI. The ribosomal binding site which was lost on pJC44 using the above restriction was re-introduced on prAC082.

### siRNA reverse transfection

Reverse siRNA transfection was performed as described in (22, 29) with minor adjustments. In brief, Genome-wide screens were performed with Dharmacon ON-TARGETplus SMART pool (pool of 4 siRNA per gene) and Qiagen Human Whole Genome siRNA Set HP GenomeWide (QU, 4 individual siRNAs for each target). Further validation screens included Ambion Silencer and Ambion Silencer Select custom libraries (with up to 6 additional siRNAs for about 1000 genes) and Sigma MISSION esiRNA libraries for 1900 genes. All screening experiments were conducted in a 384-well plate format. Each plate contained negative controls such as mock (transfection reagent only) and scrambled (non-targeting) siRNA. In addition, general siRNA controls for transfection efficiency and toxicity (e.g. Kif11, Fig. S1C) as well as positive controls (e.g. Cdc42, Rac1) that are known to have an effect on *Brucella* infection (13) were added to each plate. Based on Kif11, the average transfection efficiency reached 97.6% (91.5 – 99.99). The following specifications apply to all siRNA screens except the QU siRNA library where specifications are given in brackets. RNAiMAX in DMEM without fetal calf serum (FCS) was added to each well containing 1.6 pmol siRNA (QU: 1 pmol) or 15 ng esiRNA. Screening plates were then incubated at room temperature (RT) for 1 h. Following incubation, 500 HeLa cells were seeded per well in DMEM (FCS 10% final). Plates were incubated at 37°C and 5% CO_2_ for 72 h prior to infection. For assays in 96- and 24-well formats reverse transfections were performed in 6-well plates and subsequently reseeded in the respective plate format. On-target or control siRNAs were added to reach a final siRNA concentration of 20 nM together with RNAiMAx transfection reagent in DMEM without FCS. After 30 min of complex formation at room temperature, 110,000 HeLa cells in DMEM/10% FCS were added to each well. After 48 h transfection, cells were harvested by trypsinization and reseeded in DMEM/10%FCS (96-well plates: 2’800 cells per well; 24-well plates: 50,000 cells per well). The next day cells were infected as described hereafter. The following siRNAs used for CFU determination, colocalization experiments, and/or immunoblotting validation were purchased from Qiagen (Hilden/Germany): VPS35 (SI00760690); VPS26A (SI00760543); VPS26B (SI00631267); VPS29 (SI00760613); all star negative (0001027281); all star death kif11 (0001027299).

### Infection

For the genome-wide and confirmation screens, infections were performed in 384-well plates as described in (22, 29). In short, *B. abortus* 2308 pJC43 (*aphT::GFP* (58)) was grown in TSB medium containing 50 μg/ml kanamycin at 37°C to an OD of 0.8 - 1.1. Bacteria were then diluted in DMEM/10% FCS and added at a final MOI of 10,000. Plates were centrifuged at 400 x *g* for 20 min at 4°C to synchronize bacterial entry. After 4 h incubation at 37°C and 5% CO_2_, extracellular bacteria were killed by exchanging the infection medium by DMEM/10% FCS supplemented with 100 μg/ml gentamicin. After a total infection time of 44 h, cells were fixed with 3.7% PFA for 20 min at RT. For the entry assay, infections were performed as described in (29). In brief, transfected cells were infected with *B. abortus* 2308 pAC042.08 for 4 h after which GFP expression was induced for 4 h by the addition of Anhydrotetracycline (100 ng/ml) during the gentamicin killing of extracellular bacteria. Follow-up experiments and colocalization assays were performed according to the above-described protocol in 96-well and 24-well plates, respectively. For the colocalization assay cells were infected at a MOI of 2,000. 2 hpi cells were washed three times with DMEM/10% FCS containing gentamicin (100 μg/ml). After the indicated incubation time cells were washed three times with PBS and finally fixed for 20 min in 3.7% PFA (in PBS).

### Imaging with high-throughput microscopy

Microscopy was performed with Molecular Devices ImageXpress microscopes. MetaXpress plate acquisition wizard with no gain, 12 bit dynamic range, 9 sites per well in a 3×3 grid was used with no spacing and no overlap and laser-based focusing. DAPI channel was used for imaging nucleus, GFP for bacteria, and RFP for F-actin or dsRed of bacteria in the entry assay. Robotic plate handling was used to load and unload plates (Thermo Scientific). The objective was a 10X S Fluor with 0.45NA. The Site Autofocus was set to “All Sites” and the initial well for finding the sample was set to “First well acquired”. Z-Offset for Focus was selected manually and manual correction of the exposure time was applied to ensure a wide dynamic range with low overexposure. Images from the different siRNA screens are available upon request.

### Image analysis

Images were analyzed with the screeningBee analysis framework from BioDataAnalysis GmbH. To correct for uneven illumination inherent in wide-field microscopic imaging, an illumination correction model was computed for every plate using Cidre (60). To ensure that the Cidre-corrected image intensities fall within the range [0.0, 1.0] a linear transformation for pixel intensities was computed that maps the 0.001-quantile to 0.01 and the 0.999-quantile to 0.99 post-illumination correction. Illumination correction and intensity scaling were performed as pre-processing steps for every image prior to analysis.

To reduce the signal of *Brucella* DNA in the DAPI channel, a linear transform of the GFP channel was subtracted from the DAPI channel, with the linear transformation parameters f, o estimated in the following way: a mapping of GFP pixels to DAPI pixels was constructed so that for all intensities in the GFP images, the list of corresponding intensities in the DAPI images were recorded. For every list of DAPI intensities, only the mean intensities were retained. This creates a mapping of GFP intensities to their corresponding mean DAPI intensities. A linear regression was performed to obtain the linear parameters f, o that map the GFP channel image to the DAPI channel image. Cleaned DAPI images with a reduced *Brucella* signal were obtained by subtracting the linear transform of the GFP channel from the DAPI channel I’_DAPI_ = I_DAPI_ - (f I_GFP_ + o) as pre-processing steps for every image prior to analysis. On a random subset of 128 images, CellProfiler (61) was executed to identify Nucleus objects using “OTSU Global” segmentation in the DAPI channel, and the median, lower quartile and upper quartile segmentation thresholds of the images were retained as T_DAPI-m_, T_DAPI-lq_ and T_DAPI-uq_. On the same images, the GFP background intensity B_GFP_ was obtained as the position of the peak in the GFP intensity histogram, the dynamic range of the histogram D_GFP_ was obtained as the difference between the 99% quantile and the 1% quantile of intensity values, and the Bacteria segmentation threshold was computed as 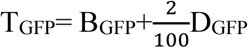. On all images, screeningBee CellProfiler was executed to perform object segmentation and measurements with the following steps: (a) Nuclei were detected as primary objects using manual threshold setting. For each plate it was manually chosen to use T_DAPI-m_, T_DAPI-lq_ or T_DAPI-uq_, depending on visual inspection of the segmentation results. Using the same threshold on all images improved site-to-site comparability. (b) Cells were detected as secondary objects around the Nuclei, with “OTSU Global” segmentation in the RFP channel. (c) Bacteria were detected as primary objects using manual threshold setting with threshold T_GFP_. Using the fixed background intensity as a reference for T_GFP_ allowed for segmenting even rather dim objects while avoiding site-to-site variability. In order to accurately measure infection scoring, a reliable method to associate pathogen colonies to individual cells is necessary. A straightforward approach is to assume that pathogen colonies must be contained within the body of the host cell. However, high cell confluence can make actin channel-based cell body segmentation inaccurate. Single microcolonies are often split into pieces that are incorrectly assigned to neighboring cells using this approach (Fig. 1B). To address this issue, we developed a novel algorithm to intelligently assign pathogen colonies to robust nucleus objects (Fig. 1C). First, inexpensive ‘bridge’ and ‘majority’ morphological operations were applied to the pathogen objects to connect broken clumps. Next, a weighted distance metric was used to measure an attraction score *a*_*N,P*_ between a pathogen *P* and individual nuclei *N* within a close proximity *d*_*prox*_. The attraction score is computed as the surface integral of the nucleus area in a continuous field emanating from the pathogen defined by an exponential function that is strongest within the microcolony itself, and decays exponentially as distance from the microcolony increases:*a*_*N,P*_ = ∑_*n*∈*N*_ *e*^−λ d_*n, p*_^, where *n*is an element (pixel) belonging to nucleus object *N*, *d*_*n,p*_ is the distance transform from the edge of microcolony *P* to *n*, and *λ* is a parameter controlling the strength of the decay. Attraction scores for all nuclei proximate to microcolony *P* are normalized such that the strongest nucleus attraction score is 1, 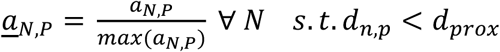. Nuclei objects with normalized attraction scores above a threshold *a*_*min*_ are associated with the pathogen microcolony. In the case that multiple nuclei are associated with the same microcolony, the microcolony is split so that each element is associated to the nearest nuclei. Large microcolonies are encouraged to split with greater ease than small microcolonies by weakening the minimum attraction score linearly according to area of the microcolony 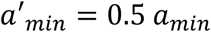 if *A*_*p*_ < *A*_*large*_ or 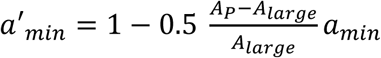 if *A*_*large*_ ≤ *A*_*p*_ ≤ 2*A*_*large*_, and 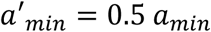 otherwise (where *A*_*p*_ is the area of the pathogen microcolony). Parameters settings *d*_*prox*_ = 45, λ = 0.2, *a*_*min*_ = 0.5, and *A*_*large*_ =8,000 were optimized by grid search on a dataset of 7,566 hand-labeled segmentations resulting in a 95.58% correct association rate.

Nucleus to pathogen microcolonies associations were aggregated. The area and integrated intensity of the pathogen objects associated to each cell and the mean intensity of the Nuclei in the GFP channel was computed as readout.

### Infection scoring for endpoint assays

Wells that contain only 32 Cells or less were excluded from infection scoring. In the remaining wells, Bacteria were filtered in a decision tree (DT) classification to exclude objects of only one-pixel area. Based on the relation of Bacteria to Nuclei, for the remaining Bacteria objects, the integrated GFP intensity was integrated over all Bacteria relating to a Cell. To reduce the impact of background intensity, an estimate for GFP background was computed using the 1% lower quantile of mean GFP intensity in the Nuclei. For every Cell, the estimated GFP background intensity was multiplied with the area of Bacteria relating to this cell, and the result was subtracted from the integrated Bacteria GFP intensity of the Cell, to arrive at a background-free estimate of “bacterial load” in each Cell. The value range for this intensity was zero for Cells with no segmented Bacteria objects, and higher than zero for all other Cells. This integrated GFP intensity was then log2-scaled, to reflect the exponential growth of replicating *Brucella*. Before log2-scaling, a small epsilon value of 2^-20^ was added to every Cell, so that the log2 value of Cells with no segmented Bacteria will not be negative infinity. The arbitrary value 2^-20^ is by a large margin smaller than the smallest actual intensity of our assays, but large enough to be used in histogram binning. For every plate, the histogram of the log2-scaled integrated cellular GFP intensity was computed (Fig. 1D) with a bin size of 0.025. The histograms were normalized to an arbitrary “virtual plate cell count” of 10^10^. To extrapolate a continuous distribution from the possibly sparse histogram, kernel density estimation (KDE) was used with a manually optimized Gaussian kernel of standard deviation 16. The histogram shows a bimodal distribution. By correlating the plate histogram distributions to selected images from the plate, we could identify that the first mode of the distribution is composed of cells with a low number of infection events ranging from single *Brucella* to small clusters (denoted as), whereas the second mode is composed of large colonies (denoted as). The two peak positions of the bimodal distribution were identified. Normal distributions G_S_ and G_L_ were fitted to the peak positions for small and large colonies, respectively. For the fitting of G_S_ and G_L_, the mean was given by the position of the peak, the height was given by the height of the peak, and the variance was optimized such that the distance between the KDE and the sum of G_S_ and G_L_ became minimal. To arrive at a binary infection scoring threshold, we identified a suitable value three standard deviations below the mean of G_L_. This threshold includes 99.8% of the events in G_L_. Cells with an integrated GFP intensity exceeding this threshold were considered true positive infections and were labelled infected. The infection score was computed as the ratio of infected Cells to the total number of Cells in the well.

### Redundant siRNA Analysis (RSA) and hit selection

Redundant siRNA Analysis RSA (25) ranks genes by iteratively assigning hypergeometric p-values to each of the multiple siRNAs targeting the same gene and picking the minimum value within a given group to represent this gene. The ranking score indicates whether the distribution of ranks corresponding to a gene is shifted towards the top, thereby aggregating the information provided by independent siRNA sequences with the same target in a robust manner. Individual siRNAs from the Qiagen library and the averages of independent replicates of the Dharmacon, Ambion, and Sigma libraries (repeated experiments with identical siRNA) were used as input. Prior to RSA analysis, siRNA targets were re-identified by searching against ENSEMBL cDNA and the REFSEQ mRNA nucleotide data, in order to ensure comparability between libraries. Cases where matching failed were excluded from this analysis. Data was further filtered removing all wells that do not pass quality control, control wells and wells where cell count was below the initially seeded cell number (500). As both up- and down hits are of interest to this analysis, RSA was run twice, once with Z-scored infection scores ranked from low to high values and once ranked oppositely. The RSA parameters were set as follows: upper and lower bound (-0.5; -2) or (-0.5;-10) on averaged z-scored infection score for down and up- hits, respectively. Bonferroni correction was applied to account for the different number of siRNAs per gene. Genes matching a Benjamini-corrected RSA P-value ≤ 0.01 with more than 3 hit wells were considered as significant and selected for further analysis.

### Infection scoring for entry assays

Wells containing only 32 Cells or less were excluded from infection scoring. In the remaining wells, Bacteria were filtered in a decision tree (DT) classification to exclude objects of only one pixel area. The remaining Bacteria were filtered in a DT classification to exclude objects of less than a manually set threshold on the upper quartile of the object intensity. The remaining Bacteria were considered true positive infections. Based on the relation of Bacteria to Nuclei, Cells were labeled infected if and only if a true positive Bacteria is related to the Cells Nuclei. The infection score was computed as the ratio of infected Cells to the total number of Cells in the well. For quantification of bacterial load in infected cells, the median of integrated GFP intensity of all true positive Bacteria was computed. The final infection readout was the product of the infection rate and bacterial load, which gives a robust approximation of the amount of intracellular bacteria (29).

### Rescue experiment

The shRNA suppression/rescue constructs for VPS35 were kind gifts from Daniel Billadeau (41). HeLa cells were seeded in a 6-well plate and transfected 4 h later with 0.9 μg of plasmid DNA using Fugene HD according to the manufacturer’s protocol. 72 h post-transfection cells were reseeded into a 96-well plate (2’800 cells / well) and infected on the following day. Cells were infected with *Brucella abortus* carrying pAC037 for 48 h. After PFA fixation and staining, cells were analyzed by image analysis. Infection scoring was performed on YFP positive cells, indicative of successful transfection.

### Determination of intracellular bacterial load by CFU determination

Infections of siRNA-transfected Hela cells were performed in 96-well plates as described above. At 6, 20, or 44 hpi, infected cells were washed with 200 µl PBS and lysed for 10 min with 0.1 % Triton X-100 / PBS. Lysed cells (6 wells per conditions) were collected in 2 ml screw-cap tubes and washed once with 1 ml PBS. Pellet was Resuspended in 1 ml PBS and subjected to 5-fold serial dilution before plating onto TSA plate. CFU were counted after 3 days growth at 37°C and normalized to the CFU obtained by the scrambled siRNA-treated cells from the matching biological replicate.

### Immunoblotting

Proteins from total cell lysates (10 - 20 µg) were separated by SDS-PAGE, transferred onto PVDF membranes (Hybond 0.2 µm, Amersham GE Life Sciences) and probed using the indicated antibodies. The secondary HRP-conjugated antibody was visualized by chemiluminescence (SeraCare developer Solution). For anti-tubulin probing, membranes were first treated with stripping buffer (Thermo Scientific), washed and reprobed. Polyclonal rabbit antibodies against VPS35 (ab97545, abcam), VPS29 (ab98929, abcam), rabbit monoclonal antibody against VPS26 (ab98929, abcam) or mouse monoclonal antibody against b-tubulin (T8328, Sigma) were used according to manufacturer’s instructions. Quantification of immunoblots was performed using ImageJ.

### Immunofluorescence for LAMP-1 co-localization

Following fixation with 3.7% PFA in PBS for 20 min, HeLa cells were incubated in PBS containing 250 mM glycine for 20 min to quench remaining aldehyde residues. Cells were then permeabilized with saponin buffer (PBS containing 0.2% saponin and 3% bovine serum albumin) for 1 h. Immunostaining was performed by incubating coverslips with saponin buffer containing antibodies against LAMP-1 (Abcam ab25630) and *Brucella abortus* LPS polyclonal rabbit serum (kind gift from Xavier De Bolle (62)) overnight in a humidified chamber at 4°C. The coverslips were then washed three times with PBS and incubated with saponin buffer containing respective fluorophore-conjugated secondary antibodies: goat anti-mouse Alexa Fluor 488 (Thermo Fisher A11029) and anti-rabbit Alexa Fluor 647 (Cell Signaling #4414) for 3 h in a humidified chamber at room temperature. The coverslips were then washed three times with PBS and mounted onto glass slides using Vectashield H-100 Antifade Mounting Medium (Vector Laboratories) and sealed with nail polish.

### Confocal microscopy for single cell data

The images were captured with the LSM-800 Confocal Microscope (Carl Zeiss) using a 63x oil objective. For each condition, 40 images were obtained at random locations across the coverslip, representing more than 50 individual cells per conditions. The images were deconvolved using Huygens software (Scientific Volume Imaging). The presence of LAMP-1 signal around the bacteria was quantified from the images by assessing the overlap between the anti-LPS and the anti-LAMP-1 staining for each individual bacterium (more than 400 per condition).

**Supporting Table S1 (SupportingTableS1-2.xls)**

List of the 223 down-hits from the genome-wide siRNA screen (RSA analysis) with associated p-value and infection score (cut-off: RSA p-value <0.01, S1_Table.xlsx).

**Supporting Table S2 (SupportingTableS1-2.xls)**

List of the 202 up-hits from the genome-wide siRNA screen (RSA analysis) with associated p-value and infection score (cut-off: RSA p-value <0.01, S2_Table.xlsx).

**Supporting Table S3 (SupportingTableS3.xls)**

Aggregated image analysis data used for Figure 3, with associated infection scores for both entry and endpoint assays. Single data points and averaged data per siRNA and per genes (as displayed in Fig. 3B) are indicated (S3_Table.xlsx).

**Supporting Table S4.** Intensity quantification from Western blot analysis of siRNA-treated HeLa cells. Presented are the averaged normalized intensities and associated standard deviation (n=3) for each VPS retromer protein upon knockdown of the indicated gene. Normalized intensities correspond to the intensity of the designated proteins divided by the intensity of the tubulin signal obtained for the same sample, on the same blot.

| siRNA | VPS35 |  | VPS29 |  | VPS26 |  |
| --- | --- | --- | --- | --- | --- | --- |
|  | AVG | SD | AVG | SD | AVG | SD |
| siVPS35 | 21.5 ± | 2.4 | 25.8 ± | 11.5 | 25.1 ± | 6.2 |
| siVPS29 | 105.8 ± | 10.4 | 6.6 ± | 5.3 | 90.2 ± | 36.8 |
| siVPS26A | 65.5 ± | 13.9 | 21.1 ± | 21.5 | 10.6 ± | 2.9 |
| siVPS26A/B | 49.5 ± | 12.9 | 23.7 ± | 31.3 | 5.5 ± | 3.3 |

## Acknowledgments

We want to thank Julia Feldmann and Dr. Sonia Borrell from the SwissTPH, Basel, Switzerland for great assistance and support with the BSL-3 environment. We are thankful to the Imaging Core Facility of the Biozentrum, Basel for excellent assistance and advice with confocal microscopy. We also thank Dr. Daniel Billadeau for sharing the VPS35 shRNA suppression/rescue constructs. Finally, we thank Dr. Simone Eicher for help with the RSA analysis and critical discussions. The genome sequence used in this research was derived from a HeLa cell line. Henrietta Lacks, and the HeLa cell line that was established from her tumor cells without her knowledge or consent in 1951, have made significant contributions to scientific progress and advances in human health. We are grateful to Henrietta Lacks, now deceased, and to her surviving family members for their contributions to biomedical research. This study was reviewed by the NIH HeLa Genome Data Access Working Group (http://acd.od.nih.gov/hlgda.htm). The genomic datasets used for analysis described in this manuscript were obtained from the database of Genotypes and Phenotypes (dbGaP) through dbGaP accession number phs000640.v1.p1. This work was supported by grants 31003A_173119 to CD from the Swiss National Science Foundation (SNSF, www.snf.ch), advanced grant 340330 to CD (FicModFun) from the European Research Council (ERC), grant 51RTP0_151029 to CD for the Research and Technology Development (RTD) project TargetInfectX in the frame of SystemsX.ch (www.systemX.ch), the Swiss Initiative for System Biology. AC and SH were fellows of the ‘Fellowship of Excellence’ International PhD program of the Biozentrum, University of Basel, Switzerland. HB benefited from the SNSF Flexibility Grant.

AC and SL performed all genome-wide siRNA screening, AC performed the entry assays, ME, KS, and AC designed and performed the image analysis for the screening and entry assays. MQ, AC, and HB performed the analysis of the genome-wide datasets. MQ and AC. analyzed the entry assay datasets. TT, SL, and HB designed and performed the VPS35 rescue experiment. TT and JS designed and performed the single cell experiment. TT, MQ, and MK. performed the CFU and WB experiments. MQ and JS analyzed the single cell data. MQ, AC and CD wrote the manuscript. CD provided strategic leadership for the project, designed experiments, and discussed data. The authors declare no competing interests.

## Supporting information legends

**Fig. S1.**
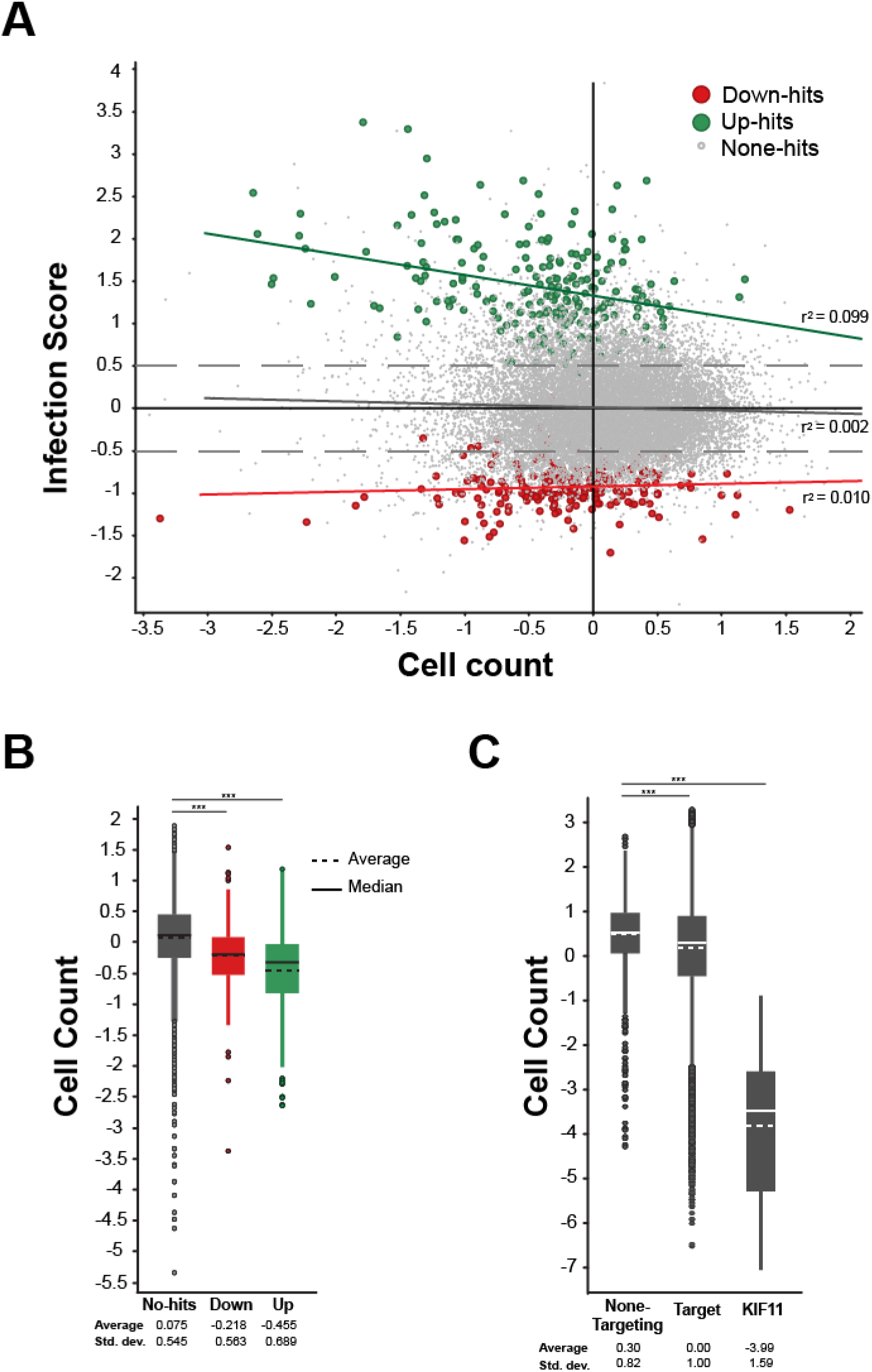
Overview of the genome-wide dataset. (**A**) Scatter plot representation of the z-scored averaged infection score plotted against the z-scored averaged cell count for all tested genes, with up- and down-hits highlighted in green and red respectively. (**A**). Linear regression between both parameters and associated r^2^ value are indicated, suggesting no direct correlation between infection score and cell count. (**B**) Box-blot representing averaged cell count score (single gene level) for different hit categories. Albeit of rather modest amplitude, a highly significant decrease in mean cell count is observed for both up- and down-hits. (**C**) Box-blot representing averaged cell count score (averaged per siRNA) for the KIF11 control compared to all targeting siRNAs or non-targeting controls. Cell number reduction in KIF11-transfected cells reached 97.6% (91.5 – 99.99). Statistical test for (B, C): Unpaired t-test with Welch’s correction, P-value < 0.0001.

**Fig. S2.**
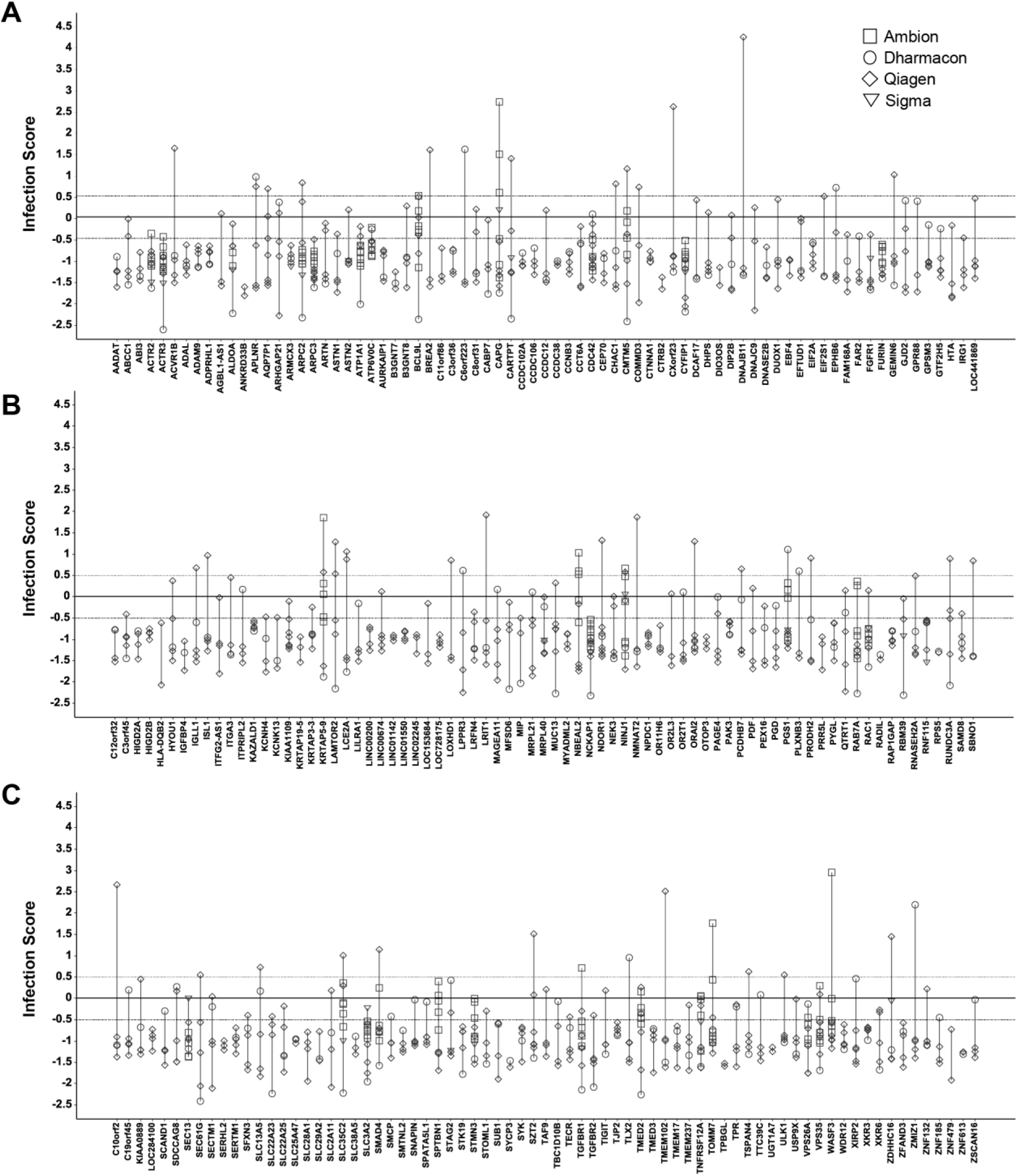
Data from single RNAi reagents for the selected 223 down-hits. (**A-C**) Averaged infection score (z-scored) and associated standard deviation for each of the 223 down-hits (RSA analysis). Displayed are the results of each tested RNAi reagents (siRNA or esiRNA), averaged by biological replicate. RNAi providers are differentiated by the shape used to indicate the data point.

**Fig. S3.**
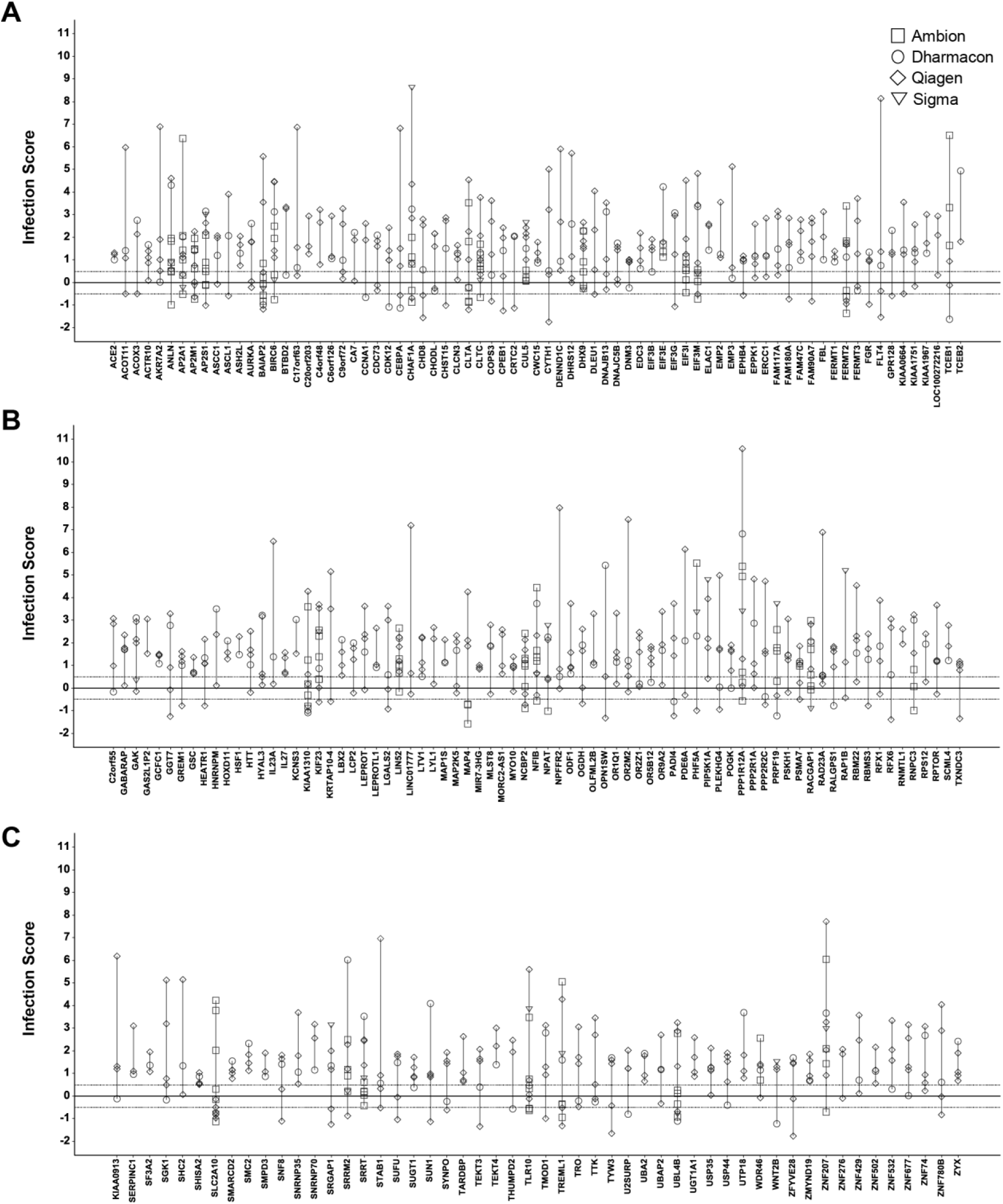
Data from single RNAi reagents for the selected 203 up-hits. (**A-C**) Averaged infection score (z-scored) and associated standard deviation for each of the 203 up-hits (RSA analysis). Displayed are the results of each tested RNAi reagents (siRNA or esiRNA), averaged by biological replicate. RNAi providers are differentiated by the shape used to indicate the data point.

